# Stac1 Regulates Sensory Stimulus Induced Escape Locomotion

**DOI:** 10.1101/2022.08.24.505176

**Authors:** Jeremy W. Linsley, Nadia Perez, I-Uen Hsu, Yuyang Yang, Naveen Jasti, Matthew Waalkes, Eric J. Horstick, John Y. Kuwada

**Author notes:** **CORRESPONDENCE**: John Y. Kuwada.

## Abstract

The *stac* family of genes are expressed by several cell types including neurons and muscles in a wide variety of animals. In vertebrates, *stac3* encodes an adaptor protein specifically expressed by skeletal muscle that regulates L-type calcium channels (CaChs) and excitation-contraction coupling. The function of Stac proteins expressed by neurons in the vertebrate CNS, however, is unclear. To better understand neuronal Stac proteins, we identified the *stac1* gene in zebrafish. *stac1* is expressed selectively in the embryonic CNS including in Kolmer-Agduhr (KA) neurons, the cerebral fluid-contacting neurons (CSF-cNs) in the spinal cord. Previously CSF-cNs in the spinal cord were implicated in locomotion by zebrafish larvae. Thus, expression of *stac1* by CSF-cNs and the regulation of CaChs by Stac3 suggest the hypothesis that Stac1 may be important for normal locomotion by zebrafish embryos. We tested to see if optogenetic activation of CSF-cNs was sufficient to induced swimming in embryos as it is in larvae. Indeed, optogenetic activation of CSF-cNs in embryos induced swimming in embryos. Next, we generated *stac1-/-* null embryos and found that both mechanosensory and noxious stimulus-induced swimming were decreased. We further found that zebrafish embryos respond more vigorously to tactile stimulation in the light compared to the dark. Interestingly, light enhancement of touch-induced swimming was eliminated in *stac1* mutants. Thus, Stac1 regulates escape locomotion in zebrafish embryos perhaps by regulating the activity of CSF-cNs.

**SIGNIFICANCE STATEMENT:** The *stac* genes are a small family of genes found in neurons and muscle in both vertebrates and invertebrates. Stac3 is a muscle protein that controls excitation-contraction coupling via regulation of L-type calcium channels and in humans a *STAC3* mutation is responsible for a congenital myopathy. The function of neural Stac proteins, however, is unknown in vertebrates. The findings of this report show that neural *stac1* is expressed by cerebral fluid-contacting neurons (CSF-cNs) in the spinal cord of zebrafish embryos and that it is necessary for normal sensory stimulus induced escape swimming. To our knowledge this is the first demonstration of a function for *stac* genes in neurons in the vertebrate nervous system.

## INTRODUCTION

In vertebrates the Stac proteins constitute a small family of proteins that uniquely contain a cysteine-rich domain (CRD) and one or two src-homology3 (SH3) domains that are expressed by several different cell types including neurons and muscles (Suzuki et al., 1996; Lein et al., 2007; Legha et al., 2010; Horstick et al., 2013; Nelson et al., 2013; Reinholt et al., 2013). In zebrafish and mice Stac3 is required in skeletal muscles for normal excitation-contraction (EC) coupling (Horstick et al., 2013; Nelson et al., 2013; Linsley et al., 2017) and in humans a missense mutation in *STAC3* is causal for Native American myopathy (Horstick et al., 2013). In mice stac3 also regulates muscle development (Gee et al., 2014: Cong et al., 2016).

In mammals *stac1* and *stac2* are expressed by subsets of neurons (Suzuki et al., 1996; Lein et al., 2007; Legha et al., 2010) but their *in vivo* neural function is unknown in any vertebrate. Recently, a *Drosophila stac* gene, *Dstac*, was identified and found to be expressed by muscles and a subset of neurons, including a specific group of clock neurons in the brain that control circadian rhythm (Hsu et al., 2018). Genetic manipulation of *Dstac* showed that Dstac in muscles was required for EC coupling (Hsu et al., 2020a) as is Stac3 in vertebrate muscles. Furthermore, Dstac in the clock neurons was necessary for normal circadian rhythm. How Dstac in the clock neurons regulate circadian rhythm is unknown but these neurons express the neuropeptide, pigment dispersing factor (PDF), that is critical for circadian rhythm (Shafer and Taghert, 2009). One way Dstac might control circadian rhythm is by regulating the activity of the clock neurons and the release of PDF.

Dstac is also expressed in the boutons of motor neurons at the neuromuscular junction in *Drosophila* larvae (Hsu et al., 2020b). In larval motor neurons manipulations of Dstac and Dmca1D, the *Drosophila* L-type calcium channel, showed that Dstac regulates Dmca1D-dependent Ca^2+^increase and the release of the neuropeptide proctolin by motor boutons and locomotion. Thus, it is possible that Stac proteins expressed by neurons in the vertebrate CNS may also regulate the activity of these neurons and behavior. Here we identified and examined Stac1, which we found is expressed by a subset of neurons in zebrafish embryos including the Kolmer-Agduhr (KA) neurons (Bernhardt et al., 1992), the cerebral fluid-contacting neurons in the spinal cord (CSF-cNs) that regulate swimming in zebrafish larvae (Wyart et al., 2009). We generated a *stac1* null mutation in zebrafish and found that it was required for normal escape swimming induced by sensory stimuli. We further discovered that light modulated sensory induced swimming and that *stac1* was necessary for this modulation by light.

## MATERIALS AND METHODS

### Animals

Zebrafish were maintained in a fish breeding facility following guidelines of the University of Michigan Animal Care and Use protocols (Zhou et al., 2008) and embryos collected and staged according to hours post-fertilization (hpf; Westerfield, 1995). All experiments were performed in compliance with the guidelines of the Institutional Animal Care and Use Committee of the University of Michigan. Transgenic fish lines used were *Tg(hb9:mgfpz)* (Flanagan-Steet et al., 2005), *Tg(1020t:gal4)* (Wyart et al., 2009) and *Tg(UAS:ChR2-mCherry)* (Scott et al., 2007). The sex of the zebrafish embryos analyzed was not determined.

### Cloning of zebrafish *stac1*, *in situ* hybridization and immunolabeling of whole-mounted embryos

To Clone Stac1, mRNA was extracted from 50 pooled wt embryos at 48 hours post fertilization (hpf) using TRIzol (Invitrogen) reagent. SuperScriptII and Oligo dT (Invitrogen) were used for reverse transcription to create a cDNA library. 5’RACE (GIBCOBRL) was performed using an intermediate 3’ primer (CCGAAGGCCCAGCTTAGCATTG) and a nested 3’ primer (CAGCTTAGCATTGGTGCCTACAATCAT) to confirm the start codon and identify the 5’ untranslated region of the stac1 locus. Full length *stac1* was cloned using the forward primer (5’-ATGATTCCGCCAGCAAACATGA-3’) and the reverse primer (5’AAATTAGTTCTGAGAAAAGCTCTGGC-3’), and was subsequently cloned into pGemT-Easy (Promega). Electrophoresis revealed a single band that was PCR purified and cloned using pGemT-easy (Promega). Sequencing was performed at the University of Michigan DNA sequencing core. The cloned plasmid was linearized with NcoI, and used as a template for antisense probe synthesis using a DIG RNA labeling kit (Roche) and Sp6 polymerase (Promega). *In situ* hybridization was conducted as previously described (Zhou et al., 2008). Anti-DIG-AP Fab fragments (Roche) were used to carry out color reactions using NBT/BCIP as a substrate (Roche). For fluorescent *stac1* and GABA double-labeling, Fast Red (Roche) was used as a substrate for *in situ* hybridization and immunolabeling was carried out with primary antibodies anti-GABA (1:500, Sigma–Aldrich), anti-GFP (1:1000, Torrey Pines Biolabs), and secondary antibody alexa488-anti-mouse/rabbit IgG (1:500, Invitrogen, Life Technologies). To assay dorsal CSF-cNs, *in situ* hybridization was performed using a *sst1.1* riboprobe (accession # NM_183070) following Molecular Instruments HCR RNA-Fish protocol for whole-mount zebrafish embryos (Schwarzkopf et al., 2021). The dorsal CSF-cNs were quantified in 3 contiguous segments within segments 9-13 of labeled embryos. Embryos were mounted in Vectashield mounting medium (Vector Laboratories) and fluorescent images were acquired with a Leica SP5 laser scanning confocal microscope or an Olympus Fluoview FV1000 microscope.

### Phylogenetic analysis of stac genes

Alignment of human STAC protein (ENSP00000273183) and STAC2 protein (ENST00000333461.6) with zebrafish stac protein (ENSDARP00000100359). was generated using Clustal Omega (Madeira et al., 2019) and visualized using ESPript (Robert and Gouet, 2014). Data for phylogenetic trees was derived from Ensembl version 69 (Villela et al., 2009). Phylogenetic trees were constructed using iTOL:Interactive tree of life (Ciccarelli et al., 2006; Letunic et al., 2019). Sequences of Stac protein paralogs used for sequence divergence tree were the following: Grass Carp (Stac1: CI_GC_24579.1, Stac3a: CI_GC_27453, Stac3b: CI_GC_28038), Zebrafish (Stac1: X P_001344470.4, Stac3: NP_001003505.1), Medaka (Stac1: XP_004082568.2, Stac2:XP_023813189.1, Stac3: XP_011475167.1), Spotted Gar (Stac1: XP015212663.1, Stac2:XP015217711.1, Stac3: XP015199515.1), Tropical Clawed Frog (Stac1: XP_031760281.1, Stac2: XP_002940986.1, Stac3: NP_001007507.1), Human (Stac1: NP_001278978.1, Stac2: NP_001338289.1, Stac3:NP_001273185.1).

### Optogenetic activation of swimming by zebrafish embryos

*gal4:1020t;UAS:EGFP* zebrafish were crossed with *UAS:ChR2-mCherry* zebrafish to obtain a *gal4:1020t; UAS:ChR2-mCherry;UAS:EGFP* line. *gal*4:1003t*;UAS:EGFP* zebrafish were crossed with *UAS:ChR2-mCherry* zebrafish to obtain a *gal4:1003t; UAS:Chr2-mCherry;UAS:EGFP* line. Progeny of crosses between were raised to 48 hpf, dechorionated enzymatically with Pronase, and then sorted for mCherry and EGFP expression under a Leica FLUO3 stereomicroscope. To assay optogenetic responses embryos were placed in a petri and illuminated with a Leica FLUO3 stereomicroscope. Each embryo was illuminated with green light for 5 sec to assay mCherry expression followed by no light for 5 sec and then cyan light for 5 sec for optogenetic activation. Cyan light activation was provided with a metal halide FLU03 lamp on a Leica FLUO3 upright microscope using an EGFP filter set (Leica) at fixed 2.5x zoom. Digital movies of light activated swimming were recorded and quantified by importing into ImageJ to measure the length of a vector representing the forward distance of tail movement in pixels after 15 seconds of cyan light illumination, though embryos that swam out of the field of vision swam further than were measured.

### Generation of *stac1* mutant

Transcription activator-like effector nucleases (TALENs) were designed to target the cysteine rich domain of Stac1. TAL3184 and TAL3185 were generated as gifts from Keith Joung (Addgene plasmid #41358 and #41359; http://n2t.net/addgene:41358; RRID:Addgene_41358). Left and Right TALEN sequences were cloned into pJDS74 and pJDS78 containing wild-type FokI nuclease domains. Cloned expression vectors were linearized, and then in vitro transcription was performed using T7 polymerase kit (Ambion). 2 nl of each TALE nuclease mRNAs (300 pg/nl) was injected into wt zebrafish embryos at the 1-cell stage. Embryos were raised to adult stage and outcrossed with the resulting F1 embryos were genotyped using sequencing of the targeted locus (zstac1F: CGAACAGCTGGTGCAACCCTAC, zstac1R: GGTGTGGGCATTGGGGTCTG). An F1 fish was isolated as a carrier for an in-del mutation in Stac1, and three more rounds of outcrosses and genotyping were performed to isolate the mutation. Next, in-crossing was performed on fish carrying the *stac1* indel mutation to create homozygous mutants. Genotyping of fin-clip DNA was used to identify homozygous stac1 mutant adults. Male and female homozygous *stac1* mutants were crossed, and mRNA was extracted and used for RT-PCR was used to assess transcription levels of *stac1* mRNA using primers overlapping Exon 1 and 2 (zstac1 Ex2F: CAGGAGTCAAAGCTGCAGAGGCTAAAG) and exons 2 and 3 (zstac1 Ex3R: CTTGCAGGCTTTGCACCGAA).

### Escape swimming assays

Embryos for behavioral analysis were enzymatically dechorionated using 2mg/ml Pronase (Protease, Type XIV, Sigma) for 20 minutes. Embryos were placed individually in the center of a testing dish (diameter: 14 cm) with a gridded sheet (grids: 6.1 × 6.1 mm) attached to the bottom of the dish and touched lightly on the tail with fine forceps (No. 5, FST, Dumont). The responses were video-recorded (Handycam, HDR-CX100, Sony). Movies were imported into ImageJ and the 30 s following the touch were manually tracked frame by frame using the ImageJ manual tracking plugin. Each embryo was touched from 1 to 3 times until a response was elicited. An embryo’s response was scored as 0 if it did not respond in 3 tail touches. The touch-induced swimming assays were performed and analyzed under blind conditions.

Swimming initiated by noxious stimuli was examined by application of the noxious agent, allyl-isothiocyanate (AITC) to embryos. AITC serves as a noxious stimulus, which induces an increase in locomotor activity in zebrafish larvae (Prober et al., 2008). Embryos (48 hpf) were individually placed in the center of a well of a 6-welled-plate filled with 2 ml fish water. For behavioral testing 20 μl of 5 mM AITC (Aldrich Chemistry, 377430) plus 1% DMSO was added to the center of the well from a micropipette (final AITC concentration = 50 μM) in experimental condition and 20 μl of fish water containing 1% DMSO added in control experiments. The behavioral response was video-recorded and analyzed with the VLC media player (0.25x to 0.5x normal speed) to quantitate the duration and the number of the forward swimming episodes during the first 2 minutes after addition of AITC or control solution.

### Light modulation of touch induced swimming

Embryos were dechorionated at approximately 24 hpf and placed in a dark incubator until 48 hpf when tail touch was applied under different light conditions. A light source (White Light Transilluminator, FB-WLT-1417, FisherScientific) was covered with two different filters in order to selectively block light of certain wavelengths. The room lights were turned off during the experiment. A red filter (Safelight filter, Kodak, No. 1A) was used to block a range of wavelengths that included λ_max_ for opn4 (470 nm, Davies et al., 2011) and TMT opsin (460 nm, Koyanagi et al., 2013) that may be important for the sensory-locomotor responses (“red light” condition). Red light did not induce swimming by itself (not shown) but allowed embryos to be video-recorded. We found that unfiltered light from the transilluminator induced swimming, but a 40% neutral density filter (“white light” condition) did not (not shown). Light was measured with a photometer. The unfiltered light intensity was approximately 1.1 × 10^15^ photons/cm^2^/sec (at 540 nm) and for the 40% neutral density filtered light the intensity was 4 × 10^14^photons/cm^2^/sec (at 540 nm). The red filtered light intensity was 1.5 × 10^14^photons/cm^2^/sec (at 650 nm). Under each condition, embryos were touched one min after placement in the central grid of a circular testing dish (diameter: 14 cm) to assay touch-induced swimming. Each embryo was touched from 1 to 3 times until a response was elicited. An embryo’s response was scored as 0 if it did not respond in 3 tail touches. The responses to touch were video recorded for 30 s.

### Statistical analysis

Statistical analyses were performed using SPSS (IBM^®^SPSS^®^Statistics). The means of the groups for each experiment were compared using Welch’s t-test when the Shapiro-Wilk test indicated that the data was normally distributed and the sample sizes of compared groups were unequal. If a given data set were not normally distributed, the non-parametric Mann-Whitney’s test was used to compare the means.

## RESULTS

### The zebrafish genome contains *stac1* but not *stac2*

Zebrafish *stac1* cDNA was cloned by RT-PCR from total RNA of 48 hpf zebrafish embryos and found that it encoded a predicted protein that was 37% identical to the zebrafish Stac3 protein and 65% identical to the human Stac1 protein with particularly high conservation in the CRD and SH3 domains (Fig. 1a). Furthermore, the predicted protein was only 45% identical to the human Stac2 protein, which was consistent with the protein corresponding to Stac1 rather than Stac2. To analyze the relationships among *stac* genes across species, phylogenetic analysis of the family of *stac* genes from sequenced vertebrate genomes was performed using Ensemble (Vilella et al., 2009). The phylogenetic tree showed that *stac* genes were present throughout the vertebrate lineage and clearly divided into *stac1*, *stac2*, and *stac3* paralog groups (Fig. 1b). A first duplication of the ancestral *stac* gene appears to have occurred ~535 million years ago (MYA) and was conserved within jawed vertebrates (Gnathostomata), at which point the muscle specific *stac3* diverged. A second duplication (~420 MYA) subdivided the neuronal version into *stac1* and *stac2.* In contrast to mammals, which all have a complement of a single *stac1*, *stac2*, and *stac3* genes, the teleost lineage showed a wide diversity in the number of *stac* genes encoded within genomes after the hypothesized whole-genome duplication of the teleost lineage (TGD) around 320 MYA (Jaillon et al., 2004) (Fig 1c). The spotted gar, whose lineage diverged from other teleosts before the TGD at around 340 MYA (Amores et al., 2011), contained a single *stac1*, *stac2*, and *stac3* genes. Surprisingly, there is no evidence of a *stac2* gene in the zebrafish sequenced genome (GRCz10; Howe et al., 2013). In contrast, other genomes within the teleost lineage such as the Japanese Medaka have two *stac2* genes, suggesting the possibility that zebrafish lost *stac2* after duplication of *stac2* genes within the teleost lineage. The only other sequenced teleost genome that appears to have lost *stac2* was the grass carp, the nearest sequenced relative of zebrafish, which diverged around 54 MYA and contained one *stac1*, no *stac2* and two *stac3* genes (Chen et al., 2017). Thus, the loss of *stac2* could be a feature of a conserved sub-branch of the Family Cyprinidae that includes zebrafish and grass carp, but excludes the common carp, which retained a copy of *stac2*. Sequencing of additional species closely related zebrafish will be necessary to see if this is the case.

**Figure 1.**
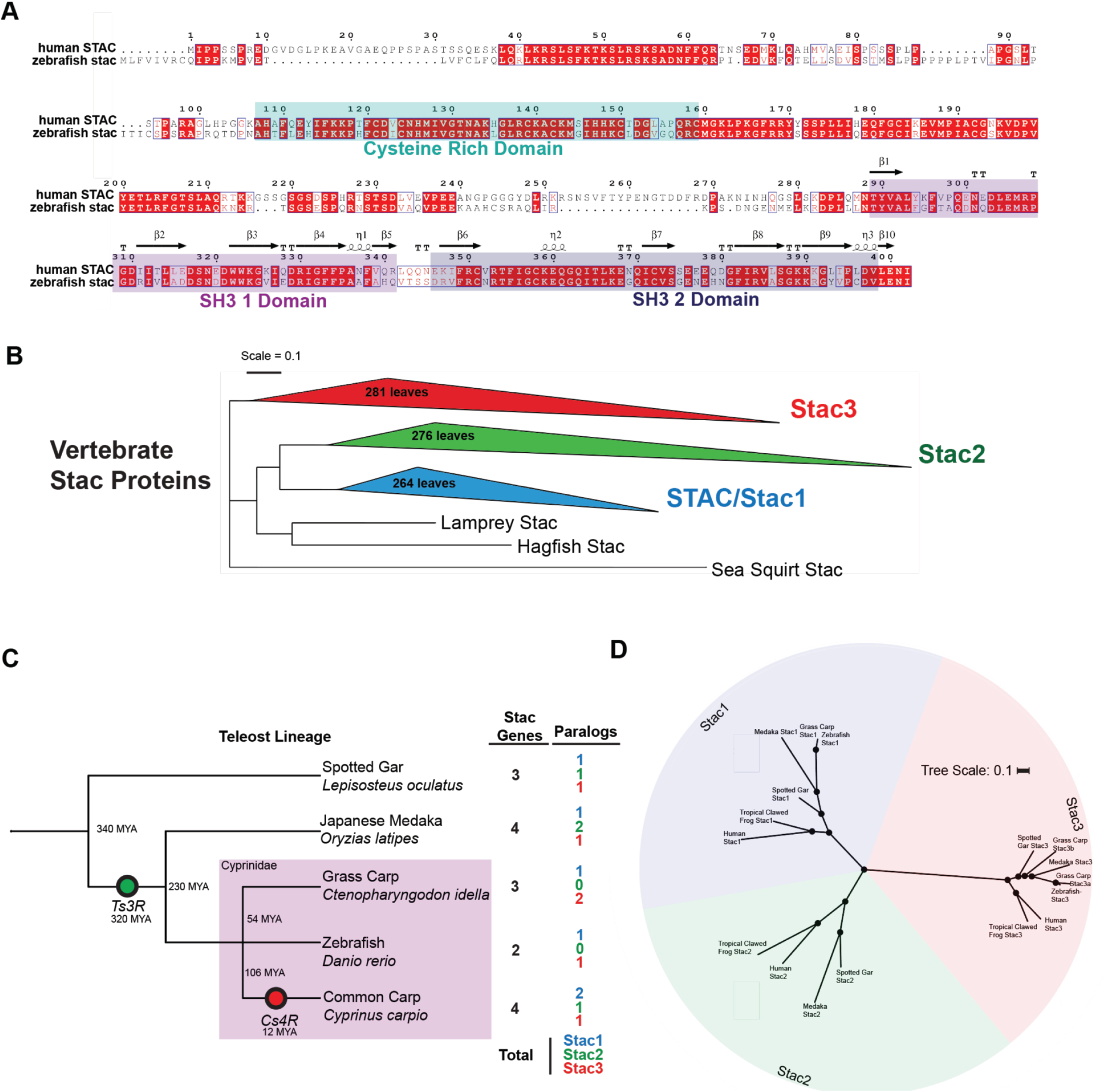
Evolution and conservation of zebrafish Stac protein. A) Amino acid alignment of Human STAC protein (ENSP 00000273183) with zebrafish stac protein (ENSDARP00000100359). Cysteine Rich Domain (turquoise) and SRC Homology 3 (SH3) domains 1 (magenta) and 2 (grey) are highlighted. Secondary structure derived from X-ray diffraction of human Stac1 SH3 domains (PDB:6B25) is displayed above sequence (Wong et al., 2017). B) Phylogeny of Stac proteins within all sequenced vertebrates (824 total) showing evolution and protein sequence separation of paralogues Stac1, Stac2 and Stac3. Species nodes were collapsed as slanted triangles and visualized within iTOLv4 (Letunic et al., 2019). C) Diversity and evolution of stac paralogs within the teleost lineage. The teleost-specific genome duplication (Ts3R, green circle), as well as carp-specific-genome duplications (Cs4R, red circle) appeared to introduce a diversity of paralog counts of Stac proteins. Approximate branch divergence dates were derived from Timetree (Kumar et al., 2017). D) Phylogenetic tree of Stac protein paralogs from human, tropical clawed frog (*Xenopus tropicalis)*, Japanese medaka, spotted gar, grass carp, and zebrafish showing grass carp and zebrafish appear to have lost stac2 genes.

### Stac1 is expressed selectively in the CNS of zebrafish embryos

To investigate the function of *stac1* we determined the expression pattern of *stac1* in zebrafish embryos. By 22 hpf *stac1* was expressed specifically by a small subset of cells in the ventral spinal cord (Fig. 2a). By 48 hpf a group of cells in the preoptic region expressed *stac1* (Fig. 2c) in addition to the ventral spinal cord cells (Fig. 2b).

**Figure 2.**
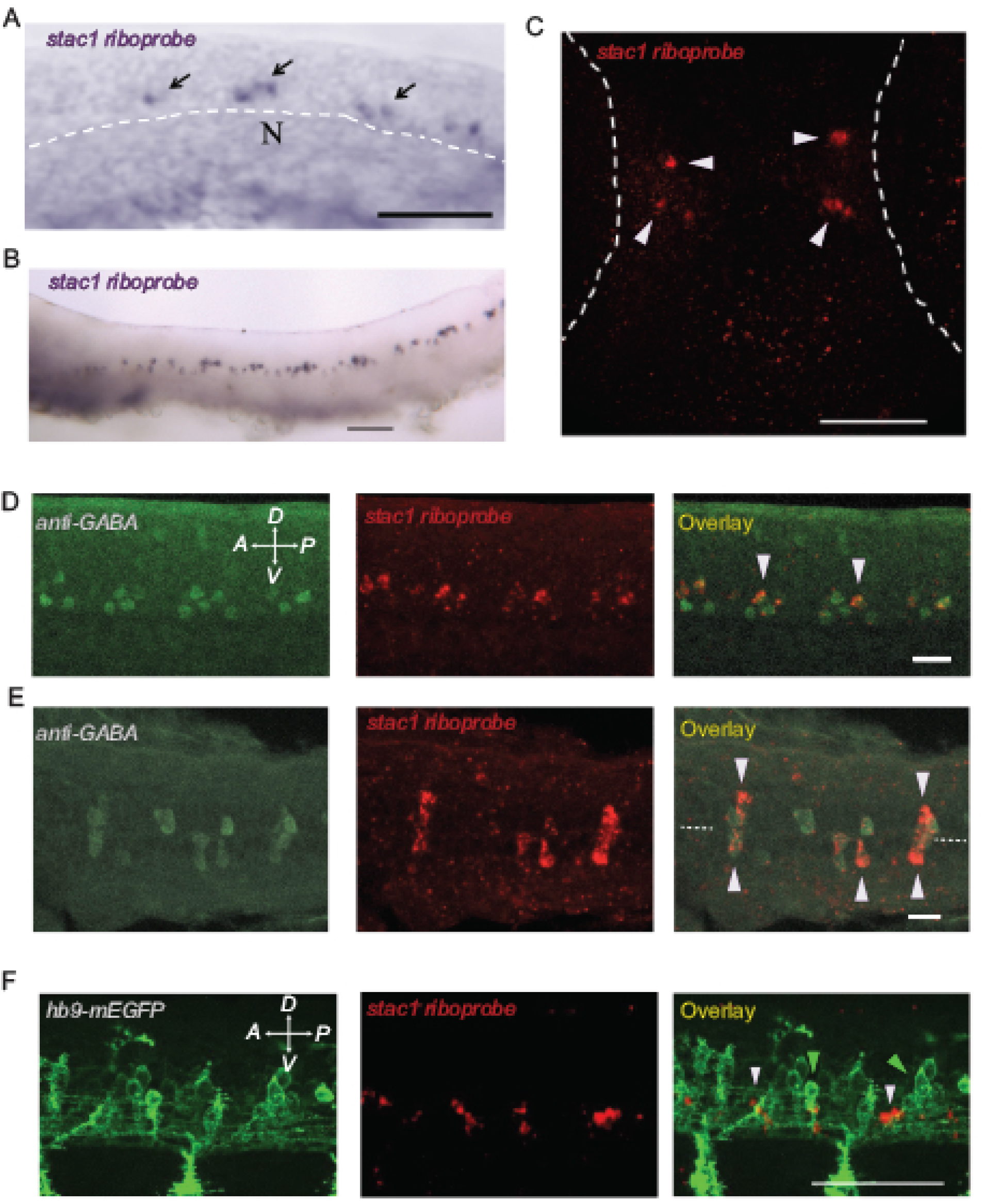
Expression of stac1 transcript in zebrafish larvae. A) Lateral view of *in situ* hybridization of WT embryos 22hpf showing *stac1* expression (black arrows) nearby the notochord (N, white dotted line indicates dorsal boundary). Scale = 25 μm. B) Lateral view of *stac1* expression at × hpf showing expression along the spinal cord. C) Expression of *stac1* at 48hpf in preoptic tectum. D) Lateral and (E) dorsal views of co-expression of *stac1* with anti-GABA labeling at 48 hpf. F)Lateral view of *stac1* expression that does not overlap with motor neuron expression of EGFP in an *hb9:*mEGFP transgenic line at 48hpf.

Since *stac1* is expressed selectively by ventral spinal cells in zebrafish embryos at a time when they exhibit simple motor behaviors, it is possible that these cells may be involved in early behavior. With this in mind, we sought to identify the *stac1* expressing cells in the ventral spinal cord. Since the embryonic ventral cord contains motor neurons (Westerfield et al., 1986), we examined whether they might express *stac1*. *In situ* hybridization with *stac1* riboprobe of *Tg*(*hb9:mgfp)* transgenic embryos (Flanagan-Steet et al., 2005), which express membrane tagged GFP in primary and secondary motor neurons, showed that motor neurons do not express *stac1* (Fig. 2f). Another set of neurons found in the early ventral spinal cord are GABA-positive Kolmer-Agduhr (KA) neurons, which are cerebral fluid-contacting neurons (CSF-cNs) in the early spinal cord that are near the ventral midline and found in two rows (Bernhardt et al., 1992; Yang et al., 2010a). Co-labeling with *stac1* riboprobe and anti-GABA showed that for the most part the more dorsal row of CSF-cNs were the *stac1*-positive cells in the early spinal cord (Fig. 2d, e).

### Optogenetic activation of cerebral fluid-contacting neurons (CSF-cNs) induces swimming by zebrafish embryos

Previously, CSF-cNs in the spinal cord of larval zebrafish were implicated in swimming (Wyart et al., 2009). Since Stac3 (Horstick et al., 2013; Nelson et al., 2013; Linsley et al., 2017) and Dstac (Hsu et al., 2018; Hsu et al., 2020) regulate the activity of muscles and some neurons, the finding that CSF-cNs express *stac1* in embryos suggested that the embryonic CSF-cNs may also be involved in swimming by embryos just as it does in larvae. However, swimming by embryos differs from that of larvae. Larval swimming involves short spurts of swimming separated by longer periods of gliding while embryonic swimming is continuous (Drapeau, et al., 2002). Since swimming differs between embryos and larvae, we investigated the role of the CSF-cNs in the spinal cord of embryos for swimming. To do this we examined whether swimming could be initiated in embryos by optogenetic activation of the CSF-cNs. We used progeny from crosses of transgenic zebrafish carrying a UAS regulated channelrhodopsin *Tg(UAS:ChR2-mCherry*) (Scott et al., 2007) with transgenic zebrafish which express Gal4 in CSF-cNs and motor neurons *Tg(1020t:gal4; UAS:EGFP*) (Wyart et al., 2009). *Tg(1020t:gal4; UAS:EGFP; UAS:ChR2-mCherry)* embryos were identified by red and green fluorescence in the spinal cord under a epi-fluorescence stereomicroscope. Indeed, 12 of 12 48 hpf embryos were found to respond to cyan light by swimming (Fig. 3a; Movie 1). One issue here is that *Tg(1020t:gal4; UAS:ChR2-mCherry)* embryos express ChR2 in motor neurons as well as CFN-cNs. One might have expected activation of spinal motor neurons would initiate uncoordinated muscle contractions. It is unknown why exposure to cyan light resulted in coordinated swimming, however, it is possible that activation of CSF-cNs may have a lower threshold compared with that of motor neurons or that ChR2 is more highly expressed in CFN-cNs than in motor neurons. To avoid this complication we examined optogenetic activation of CSF-cNs alone in embryos (*Tg(1003t:gal4; UAS:EGFP; UAS:ChR2-mCherry)*) that express Gal4 in CSF-cNs selectively (Wyart et al., 2009). In these embryos 5 of 9 responded to cyan light by swimming albeit with a delay (Fig. 3b). Thus, it appears that activation of CSF-cNs, some of which express *stac1*, was sufficient to initiate swimming in embryos prior to 3 dpf.

**Figure 3.**
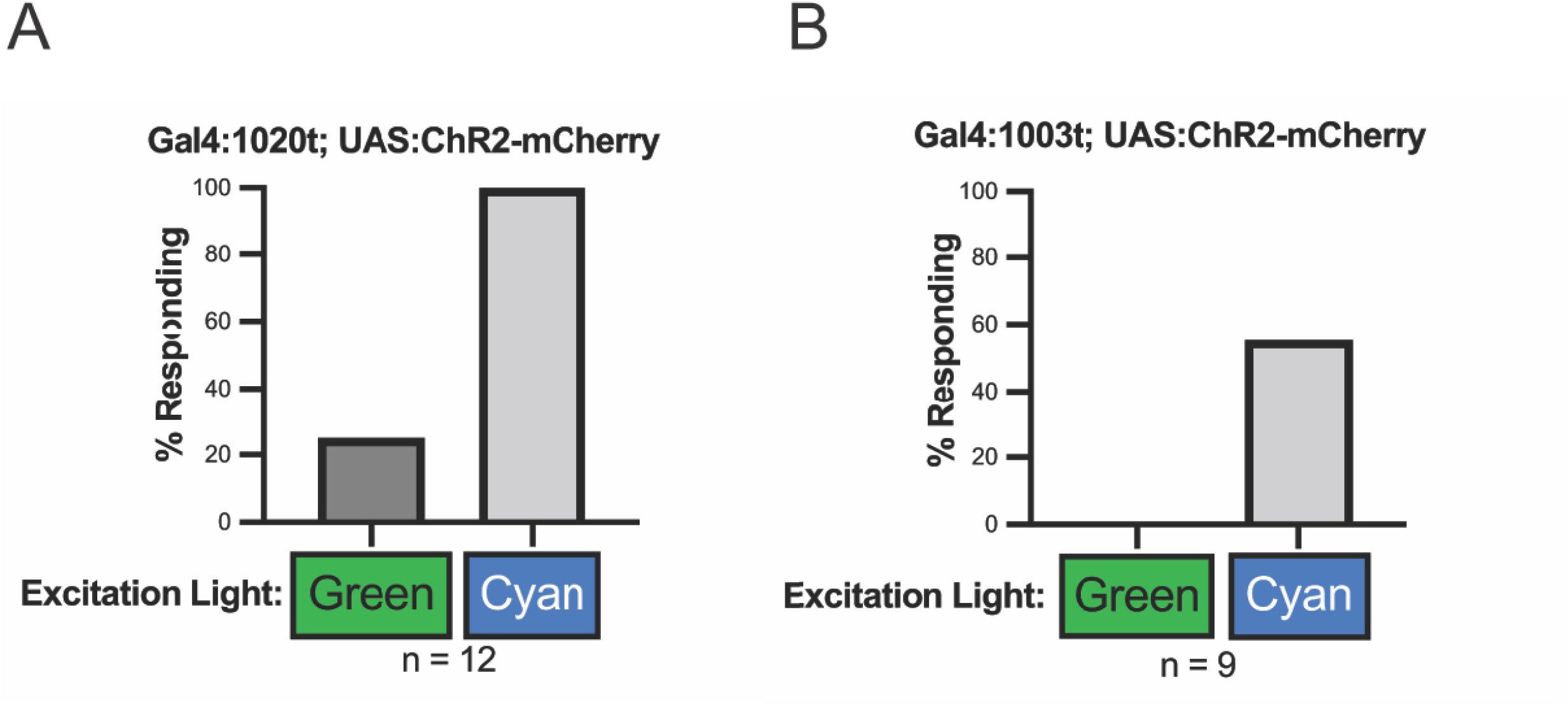
Optogenetic activation of CSF-cNs induces swimming in 48hpf zebrafish embryos. A) Quantification of the % of responding Tg:1020t:Gal4; UAS:ChR2-mCherry; UAS:EGFP embryos (48hpf) embryos to green (control) and cyan (optogenetic) light. B) Quantification of the % of responding Tg:1003t:Gal4; UAS:ChR2-mCherry; UAS:EGFP embryos (48hpf) embryos to green (control) and cyan (optogenetic) light.

### Generation of a null mutation in zebrafish *stac1*

Previously we found that a mutation in the cysteine rich domain of zebrafish Stac3 that was discovered through a mutagenesis screen leads to a null allele of the protein (Horstick et al., 2013). We looked to target a similar region of Stac1 in order to create a null mutation of zebrafish *stac1*. Transcription activator-like effector nucleases (TALENs) are an effective targeted gene knockout method in zebrafish (Sander, et al., 2011; Huang et al., 2011) that efficiently generates indel mutations (Cade, et al., 2012; Moore et al., 2012). TALEN pairs were constructed using the fast ligation-based automatable solid-phase high-throughput (FLASH) system as part of an initiative to target endogenous genes within the zebrafish genome (Reyon et al., 2012). We injected RNAs encoding TALEN pairs targeted to the CRD region of *stac1* into one-cell-stage embryos and raised them to adults. Individual TALEN-injected fish were screened by outcrossing and sequencing the *stac1* CRD locus of embryos in the F1 generation. A founder zebrafish carrying an indel mutation consisting of an 8bp deletion and 2bp insertion that resulted in a premature stop codon (Fig. 4a, b, c) was identified and isolated. Three rounds of outcrosses to wt zebrafish were used to isolate the mutation and reduce the chance of potential off-target mutations in the *stac1* mutant line. A homozygous *stac1* mutant line was then generated by mating identified male and female *stac1* mutation carriers, and identification of progeny that were homozygous *stac1* mutants was carried out by fin clip genotyping. Homozygous *stac1* mutants are healthy and viable (Movie 2), and can be bred as a stable homozygous line. The gross morphology of 48 hpf mutant embryos were indistinguishable from wt (not shown) and the dorsal CSF-cNs in the spinal cord of mutants appeared normal (Fig. 5). The dorsal CSF-cNs, express the somatostatin gene *sst1.1* (Djenoune et al., 2017) as well as *stac1* (see above) and *in situ* hybridization with a *sst1.1* riboprobe found that the distribution of dorsal CSF-cNs of 48 hpf embryos was normal in mutants. Thus, the *stac1* mutation appears not to lead to any obvious defect in the dorsal CSF-cNs in the embryonic spinal cord nor the expression of *sst1.1* by the dorsal CSF-cNs. Furthermore, RT-PCR of mRNA extracted from homozygous *stac1* mutants showed no *stac1* transcripts, suggesting that the indel mutation induced non-sense mediated decay and homozygous *stac1* mutants are null for *stac1* (Fig. 4d, e).

**Figure 4.**
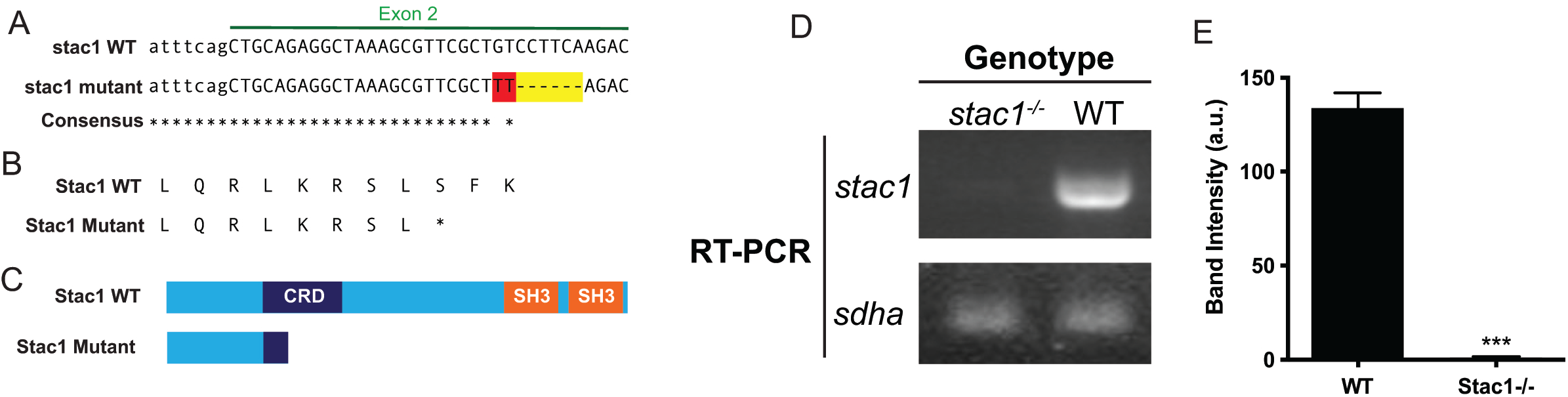
Generation of a *stac1* KO line with TALEN. A) Sequence of TALEN-induced in-del mutation site consisting of a gain of two nucleotides (red) and deletion of eight nucleotides (red and yellow) within *stac1* exon 3. B) The *stac1* indel mutation results in the introduction of an early stop codon without any additional amino acids introduced. C) A comparison of the resulting putative truncated Stac1 protein as a result of the TALEN-induced mutation to the full length WT Stac1. D) Ethidium bromide stained agarose gel of RT-PCR for *stac1* transcript from pooled *stac1-/-* and WT embryo lysate. E) Quantification of the band intensities from RT-PCR from *stac1-/-* and WT embryo lysates.

**Figure 5.**
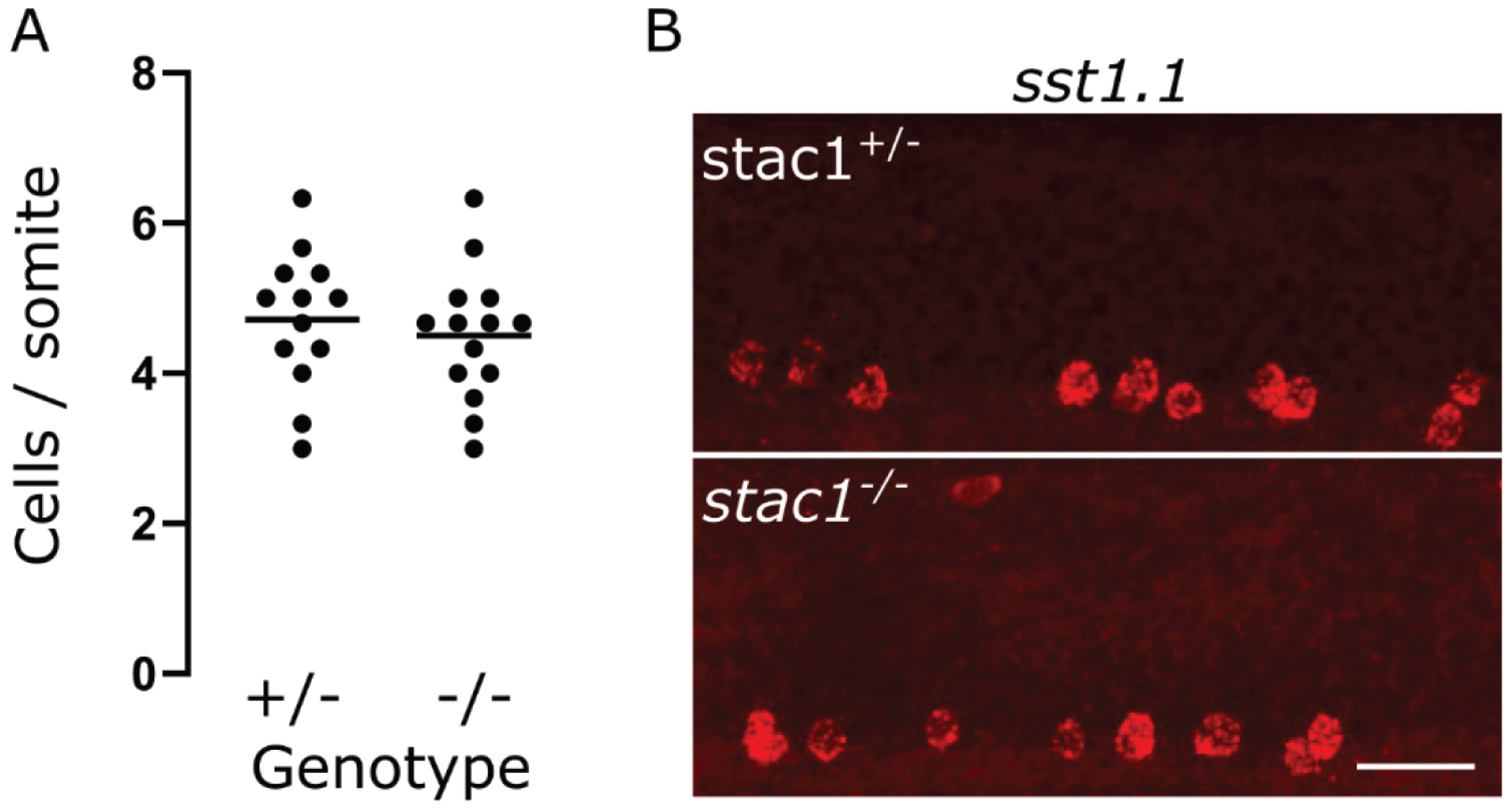
Dorsal spinal CSF-CN neurons are normal in *stac1-/-* mutant embryos (48 hpf). Dorsal spinal CSF-CNs were assayed for expression of *sst1.1*. A) Average number of *sst1.1* positive neurons per somite in *stac1+ /-* (n= 13) and *stac1-/-* (n= 14) embryos. No difference was observed (2-tailed t-test, p= 0.54). Circles are individual embryo averages across 3 somites and line denotes population average. B) Representative images showing *sst1.1* positive CSF-cNs in mid-body segments of *stac1+ /-* (top) and *stac1-/-* (bottom). Scale bar, 40μm.

### *stac1* is required for sensory stimulus evoked escape swimming

The finding that dorsal CSF-cNs in the embryonic spinal cord express *stac1* and activation of CSF-cNs in the spinal cord is sufficient to induce swimming by zebrafish embryos suggests the possibility that *stac1* in dorsal CSF-cNs might regulate swimming by embryos. We examined this hypothesis by assaying swimming initiated by sensory stimulation in *stac1* null embryos. Tactile stimulation initiated robust swimming in 48 hpf wt embryos, but touch-induced swimming was diminished in *stac1* null embryos (wt, n= 37, *stac1-/-*, n= 38, p= 0.0004 Welch’s t test; Fig. 6a). Thus, stac1 is required for normal escape swimming by zebrafish embryos initiated by tactile stimulation.

**Figure 6.**
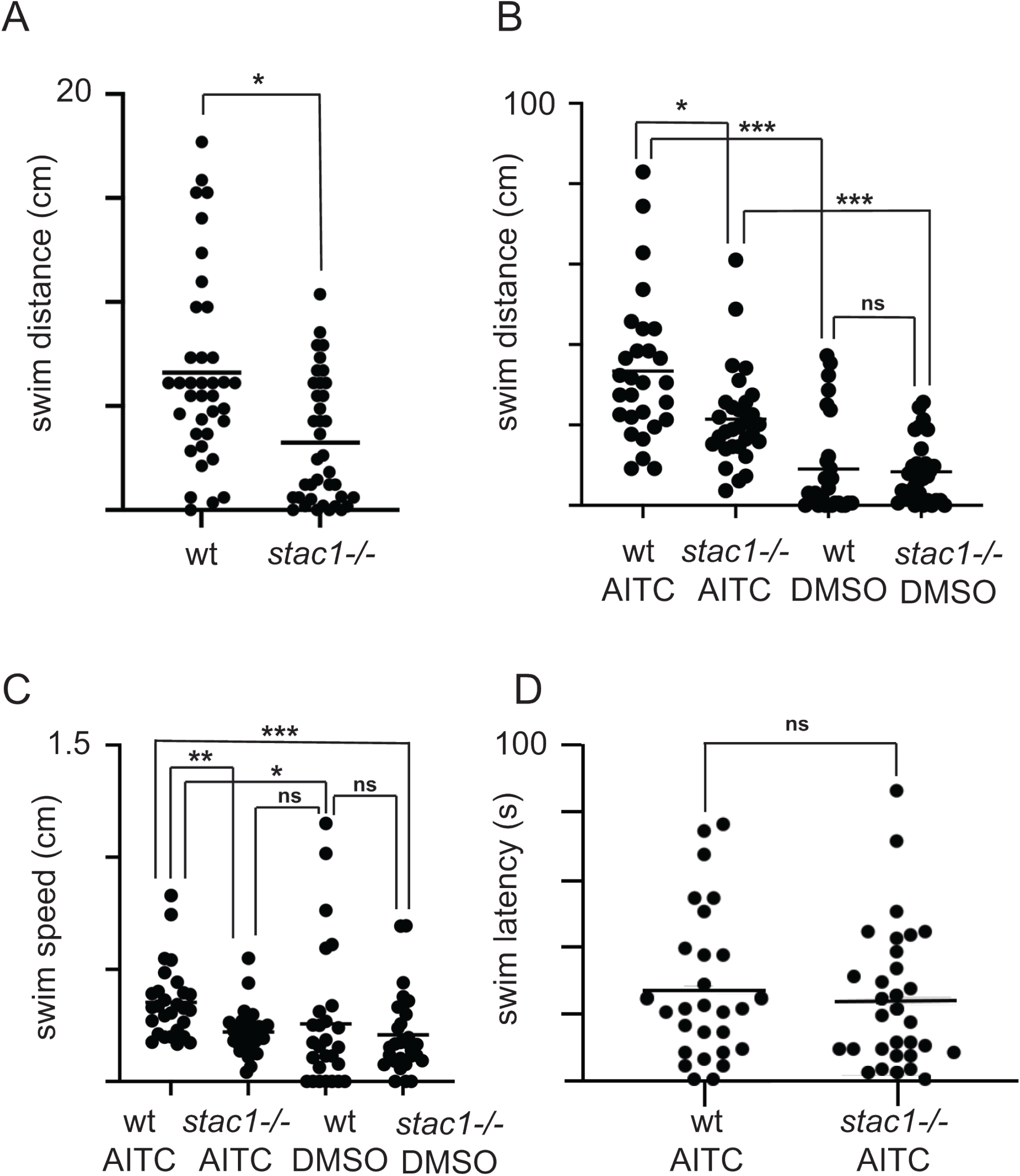
stac1^−/−^embryos (48 hpf) swim less in response to sensory stimulation. A) Swim distance is decreased in *stac1*^−/−^ embryos. Welch’s t test: wt= 37, *stac1*^−/−^= 38; p= 0.0004 (*). B) wt embryos swim in response to application of AITC compared with DMSO control and *stac1*^−/−^embryos swim less than wt. Mann-Whitney test: wt/AITC= 28, wt/DMSO= 27, *stac1*^−/−^/AITC= 30, *stac1*^−/−^/DMSO= 30. C) wt embryos swim faster with AITC compared with DMSO control and *stac1*^−/−^ embryos swim slower than wt. D) The latency of response to AITC by wt and stac1−/− embryos are comparable. In all panels B-D) Mann-Whitney tests were done with (*), (* *) and (* * *) denoting p= 0.0025, p= 0.0002 and p= 0.0001, respectively.

To examine whether the effect of the *stac1* mutation was restricted to mechanosensory stimuli induced swimming, we examined whether a potential noxious stimulus, allyl-isothiocyanate (AITC) that excites TRPA1 and induces escape swimming by zebrafish larvae (Prober et al., 2008) might also initiate swimming in zebrafish embryos. Indeed, 48 hpf wt embryos responded by swimming further when exposed to AITC (n= 28) compared with the control substance, DMSO (n= 27) (p= 0.0001, Mann-Whitney test; Fig. 6b). *stac1* null embryos also responded by swimming farther in response to AITC (n= 30) compared to DMSO (n= 30) (p= 0.0001 Mann-Whitney test) but swam less than wt embryos (p<0.0025 Mann-Whitney test). Swimming initiated by DMSO in *stac1* null embryos (n= 30) were comparable to that initiated by DMSO in wt embryos (n= 27) (p= ns, Mann-Whitney test). Furthermore, the speed of swimming initiated by AITC was higher in wt embryos (n= 28) compared with that of *stac1* null embryos (n= 30) (p= 0.0002, Mann-Whitney test) with the swimming speed initiated by AITC in *stac1* nulls (n= 30) comparable to the swimming speed initiated by DMSO in *stac1* nulls (n= 30) (p= ns, Mann-Whitney test; (Fig. 6c). Thus, wt embryos swam faster and longer than *stac1* nulls. Interestingly, the latency of response of embryos to application of AITC was similar in wt and *stac1* nulls (Fig. 6d; p= ns, Mann-Whitney test). This lack of difference in latencies is consistent with the idea that the responsiveness of *stac1* nulls to AITC was not affected by the mutation, but the amplitude of the response was. Thus, *stac1* is required for normal sensory stimulus evoked swimming by zebrafish embryos.

### Escape swimming is modulated by light and light modulation is dependent on *stac1*

The responsiveness of zebrafish embryos to sensory stimulation may be dependent on its environment. One important environmental factor might be ambient light levels so we examined whether sensory stimuli evoked swimming by embryos was influenced by light. To do this a single 48 hpf embryo in a large petri dish was placed over a light illuminator covered with a 40% neutral density filter or a red filter to modulate light levels in a dark room. A tactile stimulus was applied to the embryo and their response was video-recorded. We found that unfiltered light from the transilluminator by itself induced swimming, but a 40% neutral density filter (white light condition) did not (not shown). Illumination through the red filter (Safelight filter, Kodak, No. 1A) (red light condition) also did not induce swimming by embryos (not shown), but allowed embryos to be video-recorded in a dark room. The visual system is not functional until 68-79 hpf in zebrafish (Easter & Nicola, 1996), but the embryonic CNS contains nonretinal brain opsins (Kojima et al., 2008; Fernandes et al., 2012), which could mediate responses to light by embryos. The red filter blocked a range of wavelengths which included λ_max_ for the deep brain opsins, opn4 (470 nm, Davies et al., 2011) and TMT (460 nm, Koyanagi et al., 2013). We chose to block these opsins since they might potentially mediate light dependent modulation of behavior of zebrafish embryos. We found that touch induced a longer swimming response by wt 48 hpf embryos in the white light condition (n= 37) compared with the red light condition (n= 35) (p= 0.0029, Mann-Whitney test; Fig. 7). Thus, light appears to enhance touch induced swimming by wt embryos. However, the distances *stac1* null embryos swam following touch in the white light condition (n= 38) were comparable to that in the red light condition (n= 36) (p= ns, Mann-Whitney test) and significantly shorter than wt embryos in the white light condition (p= 0.0003, Mann-Whitney test). Touched induced swimming by *stac1* mutants was similar to that of wt under the red light condition. Thus, the *stac1* null mutation appears to eliminate light enhancement of touch induced swimming. These results are consistent with the idea that embryos respond to non-visual system mediated light stimulation, perhaps by deep brain opsins, to modulate touch evoked swimming and *stac1* is required for this modulation.

**Figure 7.**
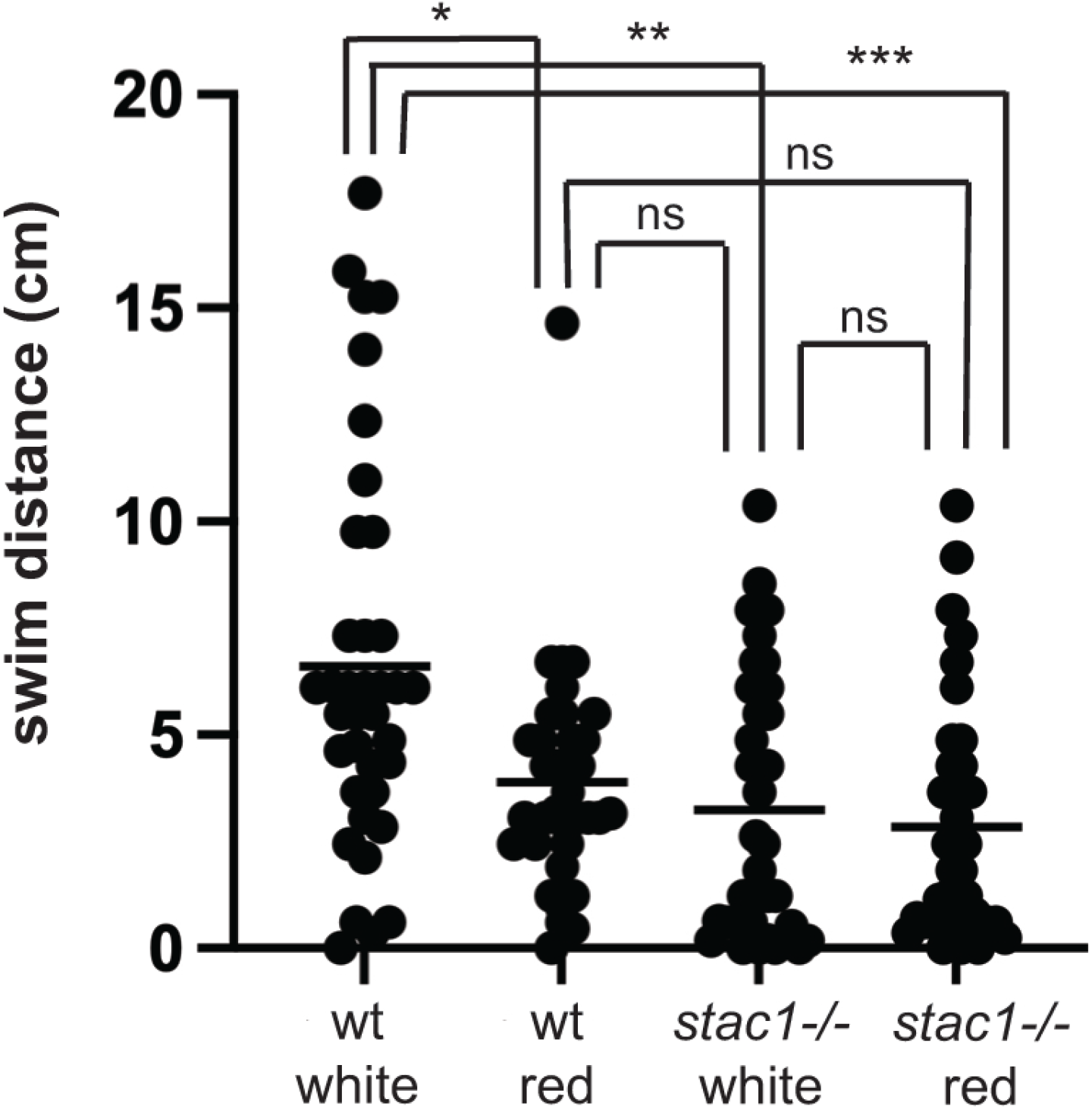
Ambient white light enhances touch induced swimming of wt embryos (48 hpf) but not *stac1*^−/−^. Mann Whitney test: wt/white light n= 37, wt/red light n= 35, *stac1*^−/−^/white light n= 36, *stac1*^−/−^ red light n= 36; p<0.003 (*), p<0.0004 (* *), p<0.0001 (* * *).

## DISCUSSION

This study found that the zebrafish genome appears to contain a single *stac1* and no *stac2* gene and that *stac1* is expressed selectively by dorsal CSF-cNs in the embryonic spinal cord. Optogenetic activation of spinal CSF-cNs in zebrafish embryos induces swimming and a null mutation in *stac1* results in decreased locomotion induced by sensory stimulation. Furthermore, escape swimming is modulated by ambient light levels and light modulation is dependent on *stac1*. This is, to our knowledge, the 1^st^ demonstration of a neuronal function for a stac gene in the vertebrate nervous system.

How *stac1* regulates escape swimming by zebrafish embryos is unknown. In *Drosophila* larvae, Dstac controls locomotion by regulating L-type calcium channels in motor neurons. Dstac is necessary for normal activity dependent increases in Ca^2+^and release of neuropeptides by motor boutons at neuromuscular junctions (Hsu et al., 2020b). Analogous to *Drosophila* Dstac it is possible that Stac1 might also regulate L-type CaChs and release of neuropeptides by the CSF-cNs in the spinal cord of zebrafish embryos. L-type CaCh are widely expressed by neurons in numerous regions of the mammalian brain (Vacher et al, 2008; Schlick et al, 2010) and the spinal cord (Jiang et al., 1999). In zebrafish embryos L-type CaChs are also widely expressed in the CNS (Sanhuenza et al, 2009). Furthermore, the spinal CSF-cNs in lamprey express somatostatin (*sst1.1*) (Jalalvand et al., 2014) and in larval zebrafish CSF-cNs contain dense-core vesicles with the dorsal CSF-cNs expressing *sst1.1* (Djenoune et al., 2017). Thus, the dorsal CSF-cNs in zebrafish embryos express both *stac1* and *sst1.1* and likely also L-type CaChs. This expression pattern is consistent with the hypothesis that Stac1 regulates the release of somatostatin by the dorsal CSF-cNs in zebrafish embryos.

Do Stac proteins regulate the release of all neuropeptides? Annotated databases of sequenced genomes (http://www.neuropeptides.nl/tabel%20neuropeptides%20linked.htm; http://isyslab.info/NeuroPep) predict that there are hundreds of neuropeptides in the brain. Given the large variation in time scales of release, sites of release and cellular actions of different neuropeptides, it would not be surprising if a number of different mechanisms regulate the release of neuropeptides. In fact, the ventral spinal CSF-cNs express the UII neuropeptides (urp1 and urp2) (Quan et al, 2015) but do not express *stac1* suggesting that Stac1 is not involved in potential release of these neuropeptides.

Since a deficiency in *stac1* results in defects in locomotion by zebrafish embryos, does somatostatin regulate locomotion? In mammals somatostatin in various motor regions of the brain such as the striatum, globus pallidus, ventral pallidum and the nucleus accumbens appears to regulate locomotion (Martin-Iverson et al., 1986; Raynor et al., 1993; Marazioti et al., 2005; Marazioti et al., 2008). In lampreys application of somatostatin to the spinal cord decreases the bursting frequency of spinal CSF-cNs and application of somatostatin receptor antagonist shortened the swimming cycle of CSF-cNs (Jalalvand et al., 2016). In zebrafish larvae the loss of illumination initiates light-seeking locomotion that is partially dependent on *sst1.1* presumably in deep brain photoreceptor neurons (Horstick et al., 2017). Since *sst1.1* is also expressed by dorsal spinal CSF-cNs (Djenoune et al., 2017), the release of spinal somatostatin may also contribute to light-seeking locomotion by zebrafish larvae.

We found that optogenetic activation of spinal CSF-cNs induced swimming by zebrafish embryos just as it does in larvae (Wyart et al., 2009). Since the dorsal CSF-cNs express *sst1.1* and somatostatin regulates locomotion in a variety of vertebrates, the dorsal CSF-cNs, which also express *stac1,* may be important for regulating swimming by zebrafish embryos. The finding that a deficiency in *stac1* reduces embryonic swimming is consistent with an important role for Stac1 in dorsal CSF-cNs for the regulation of stimulus induced swimming.

Our experiments also discovered that ambient light modulates the swimming response of 48 hpf zebrafish embryos with embryos swimming further in response to touch in the light compared to the dark. Since the visual system is not functional until 68-79 hpf in zebrafish (Easter & Nicola, 1996), it is unlikely that light modulation of touch induced swimming is mediated by the visual system. However, the embryonic CNS contains nonretinal brain opsins (Kojima et al., 2008; Fernanades et al., 2012) that could mediate light modulation. In fact, the red-light conditions used to test light modulation blocks the wavelengths of light that deep brain opsins, opn4 (Davies et al., 2011) and TMT (Koyanagi et al., 2013) respond to. Thus, it is possible that deep brain opsins might mediate light modulation of escape swimming by embryos. We further found that light enhancement of touch-induced swimming was eliminated in *stac1* mutants, revealing that light modulation of escape swimming was dependent on Stac1. Since stac1 is expressed by cells in the preoptic region of the brain at 48 hpf, the stac1 preoptic cells might mediate light enhancement of escape swimming.

How Stac1 regulates neurons in the CNS including the embryonic dorsal spinal CSF-cNs is unknown. Whether Stac1 also regulates L-type calcium channels in zebrafish neurons and thus activity dependent Ca^2+^increases are open questions. Furthermore, the requirement of Stac1 for the release of neuropeptides such as somatostatin by the dorsal CSF-cNs is unknown. These issues require further analysis to clarify the mechanism by which stac genes may control neural activity in the vertebrate nervous system.

## ACKNOWLEDGMENTS

We thank Keith Joung for TALENs against *stac1* and helpful suggestions regarding TALEN, and Kristin Verhey (University of Michigan) for sharing antibodies.

## FUNDING

Research was funded by grants from NIAMS (RO1 AR063056) to JYK and startup funds and a Research and Scholarship Advancement award from West Virginia University to EJH. JWL was supported by a Rackham Merit Fellowship from the University of Michigan and NIGMS (T32 GM007315). NP was supported by a Rackham Merit Fellowship from the University of Michigan and Research Supplement to Promote Diversity in Health Related Research from NIAMS (AR063056S). I-UH was supported by a Rackham International Student Fellowship, Rackham Barbour Scholarship, Rackham Predoctoral Fellowship, Rackham Research Grant and Rackham one-Term Fellowship from the University of Michigan. YY was supported by a Summer Research Fellowship from the University of Michigan and MW by the Ruby Distinguished Doctoral Fellowship at West Virginia University.

## DATA AVAILABILITY STATEMENT

The data that support the findings of this study are available from the corresponding author upon reasonable request.

## AUTHOR CONTRIBUTIONS

JWL, NP, I-UH, YY, MW EJH and JYK carried out the experiments and analyzed the data. JWL and JYK conceived the study and wrote the manuscript.

---

Movie 1. Tg:1020t:Gal4;UAS:ChR2-mCherry embryo swim in response to cyan light.

Movie 2. Mature *stac1*^−/−^ mutants are viable and swim normally: *stac1*−/− (left) and *stac1+/−* (right)

## Notes

**CONFLICT OF INTEREST**: The authors declare no competing financial interests.

### Competing Interest Statement

The authors have declared no competing interest.

### Summary of Updates

The stac family of genes are expressed by several cell types including neurons and muscles in a wide variety of animals. In vertebrates, stac3 encodes an adaptor protein specifically expressed by skeletal muscle that regulates L-type calcium channels (CaChs) and excitation-contraction coupling. The function of Stac proteins expressed by neurons in the vertebrate CNS, however, is unclear. To better understand neuronal Stac proteins, we identified the stac1 gene in zebrafish. stac1 is expressed selectively in the embryonic CNS including in Kolmer-Agduhr (KA) neurons, the cerebral fluid-contacting neurons (CSF-cNs) in the spinal cord. Previously CSF-cNs in the spinal cord were implicated in locomotion by zebrafish larvae. Thus, expression of stac1 by CSF-cNs and the regulation of CaChs by Stac3 suggest the hypothesis that Stac1 may be important for normal locomotion by zebrafish embryos. We tested to see if optogenetic activation of CSF-cNs was sufficient to induced swimming in embryos as it is in larvae. Indeed, optogenetic activation of CSF-cNs in embryos induced swimming in embryos. Next, we generated stac1-/- null embryos and found that both mechanosensory and noxious stimulus-induced swimming were decreased. We further found that zebrafish embryos respond more vigorously to tactile stimulation in the light compared to the dark. Interestingly, light enhancement of touch-induced swimming was eliminated in stac1 mutants. Thus, Stac1 regulates escape locomotion in zebrafish embryos perhaps by regulating the activity of CSF-cNs

## REFERENCES

Amores A, Catchen J, Ferrara A, Fontenot Q, Postlethwait JH (2011) Genome evolution and meiotic maps by massively parallel DNA sequencing: Spotted Gar, an outgroup for the teleost genome duplication. Genetics, 188:799–808.

Bernhardt RR, Chitnis AB, Lindamer L, Kuwada JY (1990) Identification of spinal cord neurons in embryonic and larval zebrafish. J Comp Neurol, 302:603–616.

Bernhardt RR, Patel CK, Wilson SW, Kuwada JY (1992) Axonal trajectories and distribution of GABAergic spinal neurons in wildtype and mutant zebrafish lacking floor plate cells. J Comp Neurol, 326:263–272.

Cade L, Reyon D, Hwang WY, Tsai SQ, Patel S, Khayter C, Joung JK, Sander JD, Peterson RT, Yeh JJ (2012) Highly efficient generation of heritable zebrafish gene mutations using homo- and heterodimeric TALENs. Nucleic Acids Res, 40: 8001–8010. https://doi.org/10.1093/nar/gks518

Chen Y, Shi M, Zhang W, Cheng Y, Wang Y, Xia X-Q (2017) The Grass Carp genome database (GCGD): an online platform for genome features and annotations. Database (Oxford), 2017:bax051.

Ciccarelli FD, Doerk T, von Mering C, Creevey CJ, Snel B, Bork P (2006) Toward automatic reconstruction of a highly resolved tree of life. Science (New York, NY), 311(5765):1283–1287.

Cong X, Doering J, Mazala DAG, Chin ER, Grange RW, Jiang H (2016) The SH3 and cysteine-rich domain 3 (Stac3) gene is important to growth, fiber composition, and calcium release from the sarcoplasmic reticulum in postnatal skeletal muscle. Skelet Muscle, 6:17.

Davies WL, Zheng L, Hughes S, Tamai TK, Turton M, Halford S, Foster RG, Whitmore D, Hankins MW (2011) Functional diversity of melanopsins and their global expression in the teleost retina. Cell Mol Life Sci, 68:4115–4132.

Djenoune L, Desban L, Gomez J, Sternberg JR, Prendergast A, Langui D, Quan FB, Marnas H, Auer TO, Rio J-P, Del Bene F, Bardet P-L, Wyart C (217) The dual developmental origin of spinal cerebrospinal fluid-contacting neurons gives rise to distinct functional subtypes. Sci Rep, 7(719):1–14. https://doi.org/10.1038/s41598-017-00350-1

Drapeau P, Saint-Amant L, Buss RR, Chong M, McDearmid JR, Brustein E (2002) Development of the locomotor network in zebrafish, Prog in Neurobiol, 68(2): 85–111. https://doi.org/10.1016/S0301-0082(02)00075-8.

Easter SS, Nicola GN (1996) The development of vision in the zebrafish (Danio rerio). Dev Biol, 180(2):646–63. https://doi.org/10.1006/dbio.1996.0335.

Fernandes AM, Fero K, Arrenberg AB, Bergeron SA, Driever W, Burgess HA (2012) Deep Brain Photoreceptors Control Light-Seeking Behavior in Zebrafish Larvae. Current Biol, 22(21):2042–2047.

Flanagan-Steet H, Fox MA, Meyer D, Sanes JR (2005) Neuromuscular synapses can form in vivo by incorporation of initially aneural postsynaptic specializations. Development:4471–4481.

Ge X, Zhang Y, Park S, Cong X, Gerrard DE, Jiang H (2014). Stac3 inhibits myoblast differentiation into myotubes. PLoS One, 9:e95926. doi: 10.1371/journal.pone.0095926

Horstick EJ, Linsley JW, Dowling JJ, Hauser MA, McDonald KK, Ashley-Koch A, Saint-Amant L, Satish A, Cui WC, Zhou W, Sprague SM, Franzini-Armstrong C, Hirata H, Kuwada JY (2013) Stac3 is a component of the excitation-contraction coupling machinery and mutated in Native American myopathy. Nat Comm 4:1952–1952.

Horstick EJ, Bayleyen Y, Sinclair JL, Burgess HA (2017) Search strategy is regulated by somatostatin signaling and deep brain photoreceptors in zebrafish. BMC Biol, 15(1):>4. https://doi.org/10.1186/s12915-016-0346-2

Howe K, Clark M, Torroja C, Stemple DL (2013) The zebrafish reference genome sequence and its relationship to the human genome. Nature, 496:498–503. https://doi.org/10.1038/nature12111

Hsu I-U, Linsley JW, Varineau J, Shafer OT, Kuwada JY (2018). Dstac is required for normal circadian activity rhythms in Drosophila. Chronobiol International, 35:1016–1026. https://doi.org/10.1080/07420528

Hsu, I-U, Linsley JW, Reid LE, Hume RI, Leflein A, Kuwada JY (2020a) Dstac regulates excitation-contraction coupling in Drosophila body wall muscles. Frontiers in Physiol, 11:573723. https://doi.org/10.3389/fphys.2020.57372

Hsu I-U, Linsley JW, Zhang X, Varineau JE, Berkhoudt DA, Reid LE, Lum MC, Orzel AM, Leflein A, Xu H, Collins CA, Hume RI, Levitan ES, Kuwada JY (2020b) Stac protein regulates release of neuropeptides. Proc Natl Acad Sci USA, 117(47):29914–29924. https://doi.org/10.1073/pnas.2009224117

Huang P, Xiao A, Shou M, Zhu Z, Lin S, Zhang B (2011) Heritable gene targeting in zebrafish using customized TALENs. Nat Biotechnol, 29(8):699–700. https://doi/10.1038/nbt.1939

Jaillon O, Aury JM, Brunet F. et al (2004) Genome duplication in the teleost fish Tetraodon nigroviridis reveals the early vertebrate proto-karyotype. Nature, 431:946–957. https://doi.org/10.1038/nature03025

Jalalvand E, Robertson B, Wallen P, Hill RH, Grillner S (2014) Laterally projecting cerebrospinal fluid-contacting cells in the lamprey spinal cord are of two distinct types. J Comp Neurol, 522:1753–1768.

Jalalvand E, Robertson B, Wallén P, Grillner S (2016) Ciliated neurons lining the central canal sense both fluid movement and pH through ASIC3. Nat Comm, 7:10002 https://doi.org/10.1038/ncomms10002

Jiang Z, Rempel J, Li J, Sawchuk MA, Carlin KP, Brownstone RM (1999) Development of L-type calcium channels and nifedipine-sensitive motor activity in the postnatal mouse spinal cord. Euro J Neurosci, 11:3481–3487.

Kojima D, Torii M, Fukada Y, Dowling E (2008) Differential expression of duplicated VAL-opsin genes in the developing zebrafish. J Neurochem, 104(5):1364–1371.

Koyanagi M, Takada E, Nagata T, Tsukamoto H, Terakita A= (2013) Homologs of vertebrate Opn3 potentially serve as a light sensor in nonphotoreceptive tissue. Proc Natl Acad Sci USA, 110:4998–5003.

Kumar S, Stecher G, Suleski M, Hedges SB (2017) TimeTree: A resource for timelines, timetrees, and divergence times. Mol Biol and Evol, 34(7):1812–1819.

Legha W, Gaillard S, Gascon E, Malapert P, Hocine M, Alonso S, Moqrich A (2010) stac1 and stac2 genes define discrete and distinct subsets of dorsal root ganglia neurons. Gene Expr Patterns, 10:368–375.

Lein ES, Hawrylycz MJ, Ao N, Ayres M, Bensinger A, Bernard A, Boe AF, Boguski MS, Brockway KS, Byrnes EJ, et al (2007). Genome-wide atlas of gene expression in the adult mouse brain. Nature, 445:168–76.

Letunic I, Bork P (2019) Interactive tree of life (iTOL) v4: recent updates and new developments. Nucleic Acids Res, 47(W1):W256–W259.

Linsley JW, Hsu I-U, Groom L, Yarotskyy V, Lavorato M, Horstick EJ, Linsley D, Wang W, Franzini-Armstrong C, Dirksen RT, Kuwada JY (2017) Congenital myopathy results from misregulation of a muscle Ca2+ channel by mutant Stac3. Proc Natl Acad Sci USA. https://doi.org/10.1073/pnas.1619238114

Madeira F, Park YM, Lee., et al. The EMBL-EBI search and sequence analysis tools APIs in 2019. Nucleic Acids Res, 47(W1):W636–W641.

Marazioti A, Kastellakis A, Antoniou K, Papasava D, Thermos K (2005) Somatostatin receptors in the ventral pallidum/substantia innominata modulate rat locomotor activity. Psychopharm, 181:319–26.

Marazioti A, Pothitos P,Papadopoulou-Daifoti Z, Spyraki C, Thermos K (2008). Activation of somatostatin receptors in the globus pallidus increases rat locomotor activity and dopamine release in the striatum. Psychopharm, 201:413–22. https://doi.org/10.1007/s00213-008-1305-6

Martin-Iverson MT, Radke JM, Vincent SR (1986) The effects of cysteamine on dopamine-mediated behaviors: evidence for dopamine-somatostatin interactions in the striatum. Pharm Biochem Behavior, 24:1707–1714.

Moore FE, Reyon D, Sander JD, Martinez SA, Blackburn JS, Khayter C, Ramirez CL, Joung JK, Langena DM (2012) Improved somatic mutagenesis in zebrafish using transcription activator-like effector nucleases (TALENs). PLoS One, 7(5):e37877. https://doi.org/10.1371/journal.pone.0037877

Nelson BR, Wu F, Liu Y., Anderson DM, McAnally J, Lin W, Cannon SC, Bassel-Duby R, Olson EN (2013). Skeletal muscle-specific T-tubule protein STAC3 mediates voltage induced Ca^2+^release and contractility. Proc Natl Acad Sci USA, 110:11881–11886.

Prober DA, Zimmerman S, Myers BR, McDermott BM Jr, Kim SH, Caron S, Rihel J, Solnica-Krezel L, Julius D, Hudspeth AJ, Schier AF (2008) Zebrafish TRPA1 channels are required for chemosensation but not for thermosensation or mechanosensory hair cell function. J Neurosci, 28(40):10102–10. https://doi.org/10.1523/JNEUROSCI.2740-08.2008

Quan FB, Dubessy C, Galant S, Kenigfest NB, Djenoune L, Leprince J, Wyart C, Isabelle Lihrmann I, Hervé Tostivint H (2015) Comparative distribution and in vitro activities of the urotensin II-related peptides URP1 and URP2 in zebrafish: Evidence for their colocalization in spinal cerebrospinal fluid-contacting neurons. PLoS One. https://doi.org/10.1371/journal.pone.0119290

Raynor K, Lucki I, Reisine T (1992) Somatostatin1 receptors in the nucleus accumbens selectively mediate the stimulatory effect of somatostatin on locomotor activity in rats. J Pharm and Exp Therapeutics, 265:67–73.

Reinholt BM, Ge X, Cong X, Gerrard DE, Jiang H (2013) Stac3 is a novel regulator of skeletal muscle development in mice. PLoS One. https://doi.org/10.1371/journal.pone.0062760

Reyon D, Tsai SQ, Khayter C, Foden JA, Sander JD, Joung JK (2012) FLASH assembly of TALENs for high-throughput genome editing. Nat Biotechnol, 30(5):460–465.

Robert X, Gouet P (2014) Deciphering key features in protein structures with the new ENDscript server. Nucleic Acids Res, 42(W1):W320–W324.

Sander JD, Cade L, Khayter C, Reyon D, Peterson RT, Joung JK, Yeh, J-R. J. (2011) Targeted Gene Disruption in Somatic Zebrafish Cells Using Engineered TALENs Nat. Biotechnol, 29(8):697–8. https://doi.org/10.1038/nbt.1934

Sanhueza D, Montoya A, Sierralta J, Kukuljan M (2009) Expression of voltage-activated calcium channels in the early zebrafish embryo. Zygote (Cambridge, England), 17:131–135.

Schlick B, Flucher BE, Obermair GJ (2010) Voltage-activated calcium channel expression profiles in mouse brain and cultured hippocampal neurons. Neuroscience, 167 (3):786–798. https:doi.org/10.1016/j.neuroscience.2010.02.037

Schwarzkopf M, Liu MC, Schulte SJ, Ives R, Husain N, Choi HMT, Pierce NH (2021) Hybridization chain reaction enables a unified approach to multiplexed, quantitative, high-resolution immunohistochemistry and in situ hybridization. Development. https://doi.org/10.1242/dev.199847

Scott EK, Mason L, Arrenberg AB, et al (2007) Targeting neural circuitry in zebrafish using GAL4 enhancer trapping. Nat Methods, 4(4):323–326.

Shafer OT, Taghert PH (2009) RNA-interference knockdown of Drosophila pigment dispersing factor in neuronal subsets: the anatomical basis of a neuropeptide’s circadian functions. PLoS One, 4:e8298.

Suzuki H, Kawaii J, Taga C, Yaoi T, Hara A, Hirose K, Hayashizaki Y, Watanabe S (1996) Stac, a novel neuron-specific protein with cysteine-rich and SH3 domains. Biochem Biophys Res Commun, 229(3):902–909.

Vacher H, Mohapatra DP, Trimmer JS (2008) Localization and targeting of voltage-dependent ion channels in mammalian central neurons. Physiol Rev, 88(4):1407–1447. https://doi.org/10.1152/physrev.00002

Vilella AJ, Severin J, Ureta-Vidal A, Heng L, Durbin R, Birney E (2009) EnsemblCompara GeneTrees: Complete, duplication-aware phylogenetic trees in vertebrates. Genome Res, 19(2):327–335.

Westerfield M (1995) The zebrafish book. Eugene, OR: University of Oregon.

Wong King Yuen SM, Campiglio M, Tung CC, Flucher BE, Van Petegem F (2017) Structural insights into binding of STAC proteins to voltage-gated calcium channels. Proc Natl Acad Sci USA, 114(45):E9520–e9528.

Wyart C, Bene FD, Warp E, Scott EK, Trauner D, Baier H, Isacoff EY (2009) Optogenetic dissection of a behavioural module in the vertebrate spinal cord. Nature, 461(7262):407–410.

Yang L, Rastegar S, Strahle U (2010) Regulatory interactions specifying Koler-Agduhr interneurons. Development. https://doi.org/10.1242/dev.048470

Zhou W, Horstick EJ, Hirata H, Kuwada JY (2008) Identification and expression of voltage-gated calcium channel beta subunits in zebrafish. Dev Dyn, 237:3842–52.

